# Separable influences of reward on visual processing and choice

**DOI:** 10.1101/2020.04.07.029942

**Authors:** Alireza Soltani, Mohsen Rakhshan, Robert J Schafer, Brittany E Burrows, Tirin Moore

**Affiliations:** Department of Psychological and Brain Sciences, Dartmouth College, Hanover, NH 03755, USA; Howard Hughes Medical Institute, Stanford University School of Medicine, Stanford, CA 94305, USA; Department of Neurobiology, Stanford University School of Medicine, Stanford, CA 94305, USA

**Keywords:** value-based decision making, attention, value integration, motion-induced bias, reinforcement learning

## Abstract

Primate vision is characterized by constant, sequential processing and selection of visual targets to fixate. Although expected reward is known to influence both processing and selection of visual targets, similarities and differences between these effects remains unclear mainly because they have been measured in separate tasks. Using a novel paradigm, we simultaneously measured the effects of reward outcomes and expected reward on target selection and sensitivity to visual motion in monkeys. Monkeys freely chose between two visual targets and received a juice reward with varying probability for eye movements made to either of them. Targets were stationary apertures of drifting gratings, causing the endpoints of eye movements to these targets to be systematically biased in the direction of motion. We used this motion-induced bias as a measure of sensitivity to visual motion on each trial. We then performed different analyses to explore effects of objective and subjective reward values on choice and sensitivity to visual motion in order to find similarities and differences between reward effects on these two processes. Specifically, we used different reinforcement learning models to fit choice behavior and estimate subjective reward values based on the integration of reward outcomes over multiple trials. Moreover, to compare the effects of subjective reward value on choice and sensitivity to motion directly, we considered correlations between each of these variables and integrated reward outcomes on a wide range of timescales. We found that in addition to choice, sensitivity to visual motion was also influenced by subjective reward value, even though the motion was irrelevant for receiving reward. Unlike choice, however, sensitivity to visual motion was not affected by objective measures of reward value. Moreover, choice was determined by the difference in subjective reward values of the two options whereas sensitivity to motion was influenced by the sum of values. Finally, models that best predicted visual processing and choice used sets of estimated reward values based on different types of reward integration and timescales. Together, our results demonstrate separable influences of reward on visual processing and choice, and point to the presence of multiple brain circuits for integration of reward outcomes.

## Introduction

Primates make approximately 3-4 saccadic eye movements each second, and thus the choice of where to fixate next is our most frequently made decision. The next fixation location is determined in part by visual salience (Itti & Koch, 2000), but also by internal goals and reward expected from the foveated target (Markowitz, Shewcraft, Wong, & Pesaran, 2011; Navalpakkam, Koch, Rangel, & Perona, 2010; Schütz, Trommershäuser, & Gegenfurtner, 2012). Brain structures known to be involved in the control of saccadic eye movement have been extensively studied as a means of understanding the neural basis of decision-making (Glimcher, 2003; Sugrue, Corrado, & Newsome, 2005). Interestingly, the same structures also appear to contribute to the selective processing of targeted visual stimuli that tends to accompany saccades (Squire, Noudoost, Schafer, & Moore, 2013). Thus, it is conceivable that reward outcomes and expected reward (i.e., subjective reward value) control saccadic choice and processing of targeted visual stimuli via similar mechanisms.

Our current knowledge of how reward outcomes and subjective reward value influence the processing of visual information and saccadic choice comes from separate studies using different experimental paradigms. For instance, the effects of reward on saccadic choice are studied using tasks that involve probabilistic reward outcomes (Chen & Stuphorn, 2015; Farashahi, Azab, Hayden, & Soltani, 2018; Liston & Stone, 2008; Platt & Glimcher, 1999; Strait, Blanchard, & Hayden, 2014) as well as tasks with dynamic reward schedules (Barraclough, Conroy, & Lee, 2004; Costa, Dal Monte, Lucas, Murray, & Averbeck, 2016; Donahue & Lee, 2015; Lau & Glimcher, 2007; Schütz et al., 2012; Sugrue, Corrado, & Newsome, 2004), both of which require estimation of subjective reward value. In contrast, the effects of reward on the processing of visual information have been mainly examined using tasks involving unequal expected reward outcomes without considering the subjective valuation of reward outcomes (B. A. Anderson, 2016; B. A. Anderson, Laurent, & Yantis, 2011a, 2011b; Barbaro, Peelen, & Hickey, 2017; Della Libera & Chelazzi, 2006, 2009; Hickey, Chelazzi, & Theeuwes, 2010, 2014; Hickey & Peelen, 2017; Peck, Jangraw, Suzuki, Efem, & Gottlieb, 2009; Rakhshan et al., 2020). More importantly, none of the previous studies has explored the effects of reward on choice and processing of visual information simultaneously. As a result, the relationship between these effects is currently unknown.

Understanding this relationship is important because the extent to which reward influences sensory processing could impact decision making independently of the direct effects of reward on choice. For example, in controlled decision-making paradigms or natural foraging settings, recent harvest of reward following saccade or visits to certain parts of the visual field or space could enhance processing of features of the targets that appear in those parts of space, ultimately biasing choice behavior. Such an influence of reward on sensory processing could have strong effects on choice behavior during tasks with dynamic reward schedules that require flexible integration of reward outcomes over time (Bari et al., 2019; Donahue & Lee, 2015; Farashahi, Donahue, et al., 2017; Farashahi, Rowe, Aslami, Lee, & Soltani, 2017; Lau & Glimcher, 2007; Soltani & Wang, 2006, 2008; Sugrue et al., 2004). In addition to better understanding choice behavior, elucidating the relationship between sensory and reward processing can also be used to disambiguate neural mechanisms underlying attention and reward (Hikosaka, 2007; Maunsell, 2004, 2015), and how deficits in deployment of selective attention, which is characterized by changes in sensory processing, are affected by abnormalities in reward circuits (Volkow et al., 2009).

Here, we used a novel experimental paradigm with a dynamic reward schedule to simultaneously measure the influences of reward on choice between available targets and processing of visual information of these targets. We exploited the influence of visual motion on the trajectory of saccadic eye movements (Schafer & Moore, 2007), motion-induced bias (MIB), to quantify sensitivity to visual motion as a behavioral readout of visual processing in a criterion-free manner. Using this measure in the context of a saccadic free-choice task in monkeys allowed us to simultaneously estimate how reward feedback is integrated to determine both visual processing and decision making on a trial-by-trial basis. We then used different approaches to compare the effects of objective reward value (i.e., total harvested reward, and more vs. less rewarding target based on task parameters) and subjective reward value (i.e., estimated reward values of the two targets using choice data) on decision making and visual processing. To estimate subjective reward values on each trial, we fit choice behavior using multiple reinforcement learning models to examine how animals integrated reward outcomes over time and to determine choice. Based on the literature on reward learning, the difference in subjective values should drive choice behavior. The MIB could be independent of subjective reward value or it could depend on subjective values similarly to or differently than choice. To test these alternative possibilities, we then used correlation between the MIB and estimated subjective values based on different integrations of reward feedback and on different timescales to examine similarities and differences between the effects of subjective reward value on choice and visual processing.

We found that both choice and sensitivity to visual motion were affected by reward even though visual motion was irrelevant for obtaining reward in our experiment. However, there were separable influences of reward on these two processes. First, choice was modulated both by objective and subjective reward values whereas sensitivity to visual motion was mainly influenced by subjective reward value. Second, choice was most strongly correlated with the difference in subjective values of the chosen and unchosen target whereas sensitivity to visual motion was most strongly correlated with the sum of subjective values. Finally, choice and sensitivity to visual motion were best predicted based on different types of reward integration and integration on different timescales.

## Methods

### Subjects

Two male monkeys (*Macaca mulatta*) weighing 6 kg (monkey 1), and 11 kg (monkey 2) were used as subjects in the experiment. The two monkeys completed 160 experimental sessions (74 and 86 sessions for monkeys 1 and 2, respectively) on separate days in the free-choice task for a total of 42,180 trials (10,096 and 32,084 trials for monkeys 1 and 2, respectively). Each session consisted of approximately 140 and 370 trials for monkeys 1 and 2, respectively. All surgical and behavioral procedures were approved by the Stanford University Administrative Panel on Laboratory Animal Care and the consultant veterinarian and were in accordance with National Institutes of Health and Society for Neuroscience guidelines.

### Visual stimuli

Saccade targets were drifting sinusoidal gratings within stationary, 5°–8° Gaussian apertures. Gratings had a spatial frequency of 0.5 cycle/° and Michelson contrast between 2%–8%. Target parameters and locations were held constant during an experimental session. Drift speed was 5°/s in a direction perpendicular to the saccade required to acquire the target. Targets were identical on each trial with the exception of drift direction, which was selected randomly and independently for each target.

### Experimental paradigm

After acquiring fixation on a central fixation spot, the monkey waited for a variable delay (200–600 *ms*) before the fixation spot disappeared and two targets appeared on the screen simultaneously (**Fig. 1A**). Targets appeared equidistant from the fixation spot, and diametrically opposite one another. The monkeys had to make a saccadic eye movement to one of the two targets in order to select that target and obtain a possible reward allocated to it (see Reward schedule). Both targets disappeared at the start of the eye movement. If the saccadic eye movement shifted the monkey’s gaze to within a 5–8°-diameter error window around the target within 400 *ms* of target appearance, the monkeys received a juice reward according to the variable reward schedule described below.

**Figure 1.**
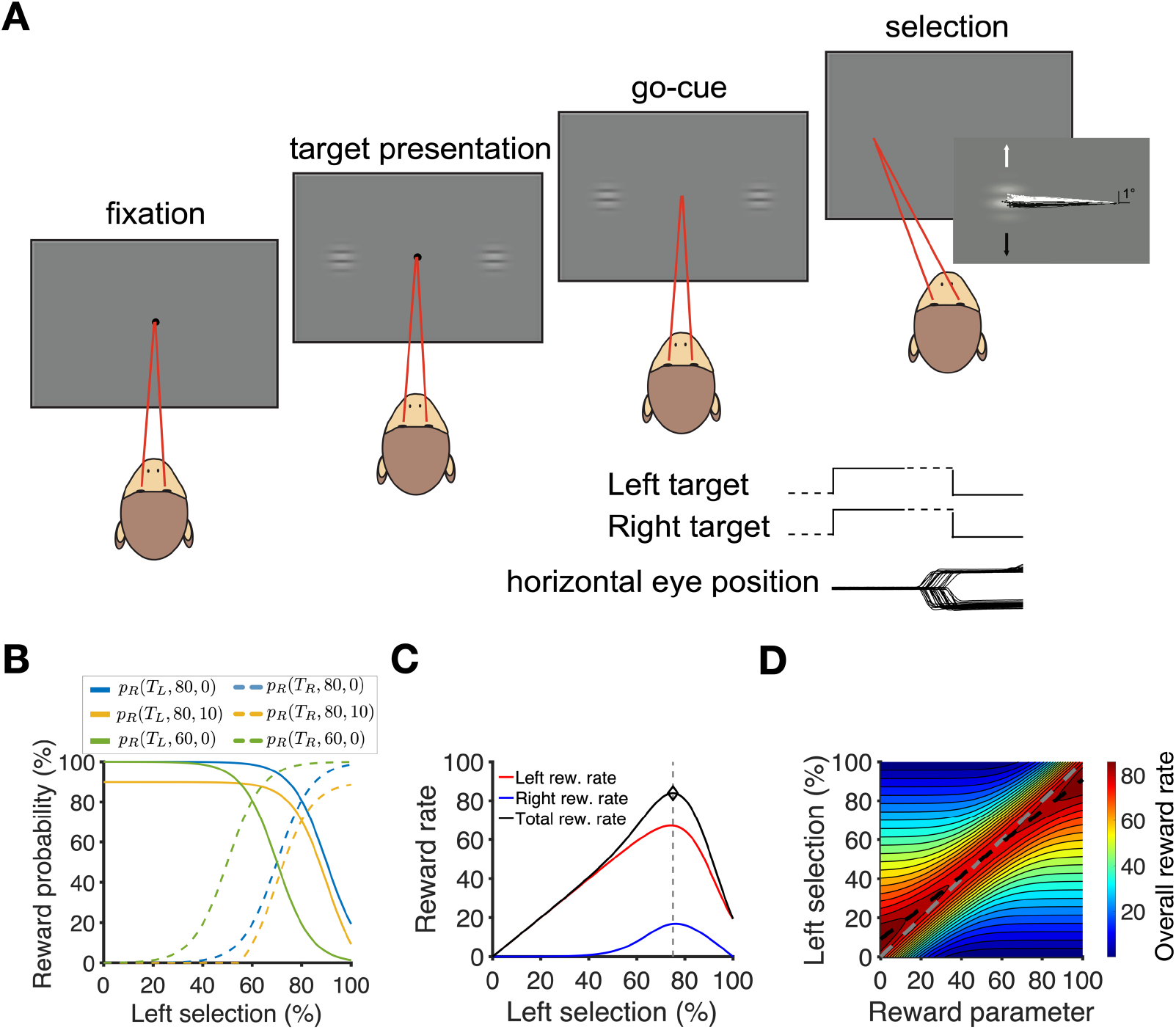
The free-choice task and reward schedule example. (**A**) Task design. On each trial, a fixation point appeared on the screen, followed by the presentation of two drifting-grating targets. The monkeys indicated their selection with a saccade. Targets were extinguished at the onset of the saccade. A juice reward was delivered on a variable schedule following the saccade. Event plots indicate the sequence of presentation of the visual targets; dashed lines denote variable time intervals. Horizontal eye position traces are from a subset of trials of an example experiment, and show selection saccades to both left target (*T_L_*, downward deflecting traces) and right target (*T_R_*, upward deflecting traces). (**B**) Examples of reward probability as a function of the percentage of left choices, separately for left and right targets (*p_R_* (*T_L_, r, x*) and *p_R_* (*T_R_, r, x*)) for different values of reward parameter *r* and penalty parameter *x* (see Eq. 1). (**C**) Plotted is the reward harvest rate on each target as a function of the percentage of *T_L_* selections, *f*(*T_L_*), for *r*=80 and *x*=0. (**D**) Total reward harvest rate as a function of reward parameter *r* and the percentage of *T_L_* selections for *x*=0. The gray dashed line shows *f*(*T_L_*) = *r* corresponding to matching behavior. The black dashed line indicates the percentage of *T_L_* selections that results in the optimal reward rate. Slight undermatching corresponds to optimal choice behavior in this task.

### Quantifying the motion-induced bias

Eye position was monitored using the scleral search coil method (Fuchs & Robinson, 1966; Judge, Richmond, & Chu, 1980) and digitized at 500 Hz. Saccades were detected using previously described methods (Schafer & Moore, 2007). Directions of drifting gratings were perpendicular to the saccade required to choose the targets. Saccades directed to drifting-grating target are displaced in the direction of visual motion, an effect previously referred to as the motion-induced bias (MIB) (Schafer & Moore, 2007). The MIB for each trial was measured as the angular deviation of the saccade vector in the direction of the chosen target’s drift, with respect to the mean saccade vector from all selections of that target within the session. This method for measuring deviation would yield approximately the same results as vertical displacement because the locations of targets were held constant throughout the session and angles were small, making angles a good approximation for the tangent of angles times the horizontal distance of the targets (vertical displacement). In order to compare MIB values across sessions with different target contrasts and locations, we used z-score values of the MIB in each session to avoid confounds due to systematic biases.

### Reward schedule

For each correct saccade, the monkey could receive a juice reward with a probability determined by a dynamic reward schedule based on the location of the foveated target (Abe & Takeuchi, 1993). More specifically, the probability of reward given a selection of the left (*T_L_*) or right (*T_R_*) target was equal to:

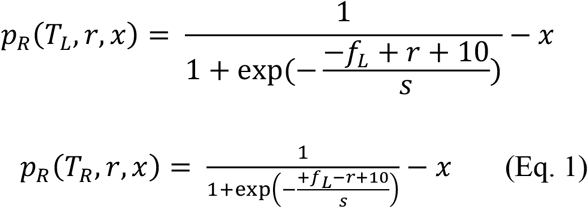

where *f_L_* is the local fraction (in percentage) of *T_L_* selections estimated using the previous 20 trials; *r* (reward parameter) is a task parameter that was fixed on a given session of the experiment and determined which option was globally more valuable (*T_L_* for *r*>50, and *T_R_* for *r*<50); *s* is another task parameter that determines the extent to which the deviation from matching (corresponding to *f_L_ =r*) results in a decrease in reward probability and was set to 7 in all experimental sessions; and *x* is a penalty parameter that reduced the global probability of a reward. Positive values of *x* decreased reward probability on saccades to both left and right targets in order to further motivate monkeys to identify and choose the more rewarding location at the time. *x* was kept constant throughout a session and was assigned to one of the following values on a fraction of sessions (reported in the parentheses in percentage): 0 (77%), 0.15 (6%), 0.30 (6%), or 0.40 (11%). Although the introduction of penalty decreased the reward probability and rate on both targets, it did not change the local choice fraction (*f_L_*) at which the optimal reward rate or matching could be achieved. Because of the penalty parameter and the structure of the reward schedule, *p_R_* (*T_L_, r, x*) and *p_R_* (*T_R_, r, x*) are not necessarily complementary. Finally, to ensure that the reward probabilities would not have negative values, any negative reward probability (based on Eq. 1) is replaced with 0.

Based on the above equations, the reward probabilities on saccades to left and right targets are equal at *f_L_ =r*, corresponding to matching behavior, which is slightly suboptimal in this task. As shown in **Fig. 1C, D**, an optimal reward rate is obtained via slight undermatching. As the value of *s* approaches zero, matching and optimal behavior become closer to each other.

### Reinforcement learning models

In our experiment, reward was assigned based on target location (left vs. right) and thus the targets’ motion directions were irrelevant for obtaining reward. Nevertheless, we considered the possibility that monkeys could incorrectly assign value to motion direction. We used various reinforcement learning (RL) models to fit choice behavior in order to determine whether monkeys attributed reward outcomes to target locations or target motions, and how they integrated these outcomes over trials to estimate subjective values and guide choice behavior. Therefore, we considered RL models that estimate subjective reward values associated with target locations as well as RL models that estimate subjective reward values associated with the motion of the two targets.

In the models based on the location of the targets (location-based RLs), the left and right targets (*T_L_* and *T_R_*) were assigned subjective values *V_L_*(*t*) and *V_R_*(*t*), respectively. In the models based on motion direction of the targets (motion-based RLs), subjective values *V_U_*(*t*) and *V_D_*(*t*) were assigned to the upward and downward motion (*T_U_* and *T_D_*), respectively. For both types of models, values were updated at the end of each trial according to different learning rules described below. In addition, we assumed that the probability of selecting *T_L_* (or *T_U_* in motion-based RLs) is a sigmoid function of the difference in subjective values as follows:

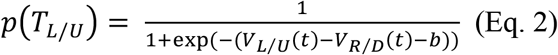

where *b* quantifies the bias in choice behavior toward the left target (or upward motion), *V_L/U_* denotes the subjective value of the left target in the location-based RL or upward motion in the motion-based RL, respectively. Similarly, *V_R/D_* denotes the subjective value of the right target in the location-based RL or downward motion in the motion-based RL, respectively.

At the end of each trial, subjective reward values of one or both targets were updated depending on the choice and reward outcome on that trial. We considered different types of learning rules for how reward outcomes are integrated over trials and grouped these learning rules depending on whether they estimate a quantity similar to *return* (average reward per selection) or *income* (average reward per trial). More specifically, on each trial, the monkeys could update subjective reward value of the chosen target only, making the estimated reward values resemble local (in time) return. Alternatively, the monkeys could update subjective reward values of both the chosen and unchosen targets, making these values resemble local income. We adopted these two methods for updating subjective reward values because previous work has shown that both local return and income can be used to achieve matching behavior (Corrado, Sugrue, Seung, & Newsome, 2005; Soltani & Wang, 2006; Sugrue et al., 2004). In addition, subjective reward values for the chosen and unchosen targets could decay similarly or differently, and monkeys could learn differently from positive (reward) and negative (no reward) outcomes. We tested all these possibilities using four different types of RL models.

In return-based RL models (RL_ret_), only the subjective value of the chosen target (in terms of location or motion direction) was updated. More specifically, if *T_L_*(*T_U_*) was selected and rewarded on trial *t*, subjective reward values were updated as the following:

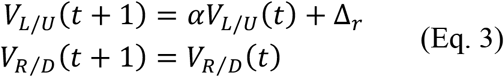

where *Δ_r_* quantifies the change in subjective reward value after a rewarded trial and *α* (0 ≤ *α* ≤ 1) is the decay rate (or discount factor) measuring how much the estimated subjective reward value from the previous trial is carried to the current trial. As a result, values of *α* closer to 1 indicates longer lasting effects of reward or integration of reward on longer timescales both of which indicate slower learning. In contrast, values of *α* closer to 0 indicate integration of reward on shorter timescales corresponding to faster learning. We note that our learning rule is not a delta rule and because of its form, (1 − *α*) in our models more closely resemble learning rate in RL models based on the delta rule. If *T_L_*(*T*_+_) was selected but not rewarded, subjective reward values of the two target locations or motion directions were updated as the following:

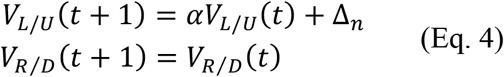

where *Δ_n_* quantifies the change in subjective reward value after a non-rewarded trial. Similar equations governed the update of subjective reward values when *T_R_*(*T_D_*) was selected. Importantly, in these models, subjective reward value of the unchosen target (in terms of location or motion) is not updated, making these models return-based.

In contrast, in all other models, subjective reward values of both chosen and unchosen targets were updated in every trial, making them income-based models. Specifically, in the RL_Inc_(1) models, the subjective value of the unchosen target decayed at a rate similar to the subjective value of the chosen target. For example, when *T_L_*(*T_U_*) was selected, the subjective values were updated as follows:

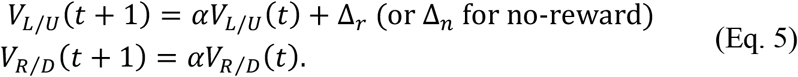

In the RL_Inc_(2) models, subjective value of chosen and unchosen targets decayed differently:

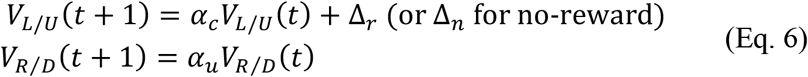

where *α_c_*, and *α_u_* are the decay rates for the chosen and unchosen targets or motion directions.

In the RL_Inc_(3) models, we updated the subjective value of unchosen target location (or unchosen motion direction) in addition to decaying the subjective values of chosen and unchosen locations:

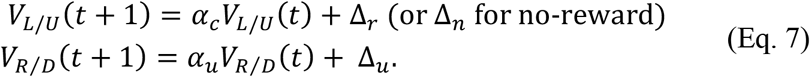

Note that the motion directions of the two targets were the same in half of the trials. This makes updating of subjective value of motion directions non-trivial in trials in which the chosen and unchosen motion directions are the same (referred to as match trials). Therefore, we tested different update rules for match trials to identify the model that best describes the monkeys’ choice behavior. Specifically, we tested two possibilities: 1) update the subjective value of motion direction that was presented on a given match trial only; 2) update the subjective values of both present and non-present motion directions but in the opposite direction. We found that the second model, in which subjective values of both motion directions were updated, provided a better fit for our data (data not shown).

Finally, we also tested hybrid RL models in which subjective values of both target locations and motion directions were updated at the end of each trial, and subsequently used to make decisions. Fitting based on these hybrid models were not significantly better than those using the RL models that consider only subjective values of target locations. Therefore, the results from these hybrid models are not presented here.

### Model fitting and comparison

We used the maximum likelihood ratio method to fit choice behavior with different RL models described above and estimated the parameters of those models. To compare the goodness-of-fit based on different models while considering the number of model parameters, we used the negative log-likelihood (-*LL*), Akaike information criterion (AIC) and Bayesian information criterion (BIC). AIC is defined as:

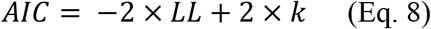

where *LL* is log-likelihood of the fit and *k* is the number of parameters in a given model.

BIC is defined as:

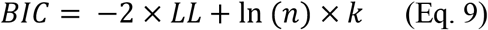

where *LL* is log-likelihood of the fit, *k* is the number of parameters in a given model, and *n* is the number of trials in a given session. We then used the best RL model in terms of predicting choice behavior to examine whether the MIB is also affected by subjective reward value similarly to or differently than choice (see below).

### Effects of subjective reward value on MIB

In order to estimate subjective reward values associated with a given target location, we used two methods of reward integration corresponding to income and return. To calculate the subjective income for a given target location on a given trial, we filtered the sequences of reward outcomes on preceding trials (excluding the current trial) using an exponential filter with a given time constant τ, assigning +1 to rewarded trials and Δ_*n*_ to non-rewarded trials if that target location was chosen and 0 if that target location was not chosen on the trial. To calculate the subjective return of a given target location, we filtered reward sequence on preceding trials (again excluding the current trial) in which that target location was chosen using an exponential filter with a given time constant τ, assigning +1 to rewarded trials and Δ_*n*_ to non-rewarded trials. Finally, we calculated the correlation between the MIB and the obtained filtered values for different values of τ and Δ_*n*_.

### Data analysis

To assess the overall performance of the monkeys, we used static and dynamic models to harvest maximum rewards. In the static model, we assumed that selection between the two target locations in a given session was a stochastic process with a fixed probability that is optimized for a given set of parameters. Replacing *f_L_* with to-be-determined probability *p*(*T_L_*) in Eq. 1, one can obtain the total average reward on the two targets, *R_tot_*, as follows:

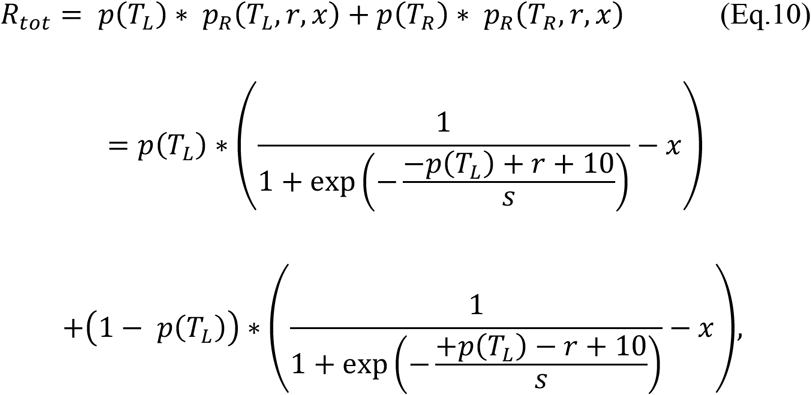

The optimal probability, *p_opt_*(*T_L_*), was then determined by maximizing *R_tot_*:

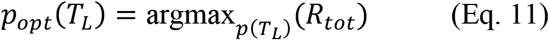

In the optimal dynamic model, we assumed that the decision maker has access to all the parameters of the reward schedule (*r, s, x*) and perfect memory of their own choices in terms of *f_L_*. Having this knowledge, the optimal decision maker could compute the probability of reward on the two options, *p_R_*(*T_L_, r, x*) and *p_R_*(*T_R_, r, x*), (using Eq. 1) and choose the option with the higher reward probability on every trial.

We also compared the monkeys’ choice behavior with the prediction of the matching law. The matching law states that the animals allocate their choices in a proportion that matches the relative reinforcement obtained by the choice options. In our experiment, this is equivalent to the relative fraction of left (respectively, right) choices to match the relative fraction of incomes on the left (respectively, right) choices. Therefore, to quantify deviations from matching, we calculated the difference between the relative fraction of choosing the more rewarding target (left when *r* >50 and right when *r* <50) and the relative fraction of the income for the more rewarding target. Negative and positive values correspond to undermatching (choosing the better option less frequently than the relative reinforcement) and overmatching, respectively.

## Results

We trained two monkeys to freely select between two visual targets via saccadic eye movement (**Fig. 1A**). Saccades to each target resulted in delivery of a fixed amount of juice reward with a varying probability. Targets were stationary apertures of drifting gratings and the reward probability was determined based on the location of the grating targets independently of the direction of visual motion contained within the gratings. More specifically, on a given trial, probabilities of reward on the left and right targets were determined by the reward parameter (*r*) and the choice history on the preceding 20 trials (Eq. 1; **Fig. 1B**). Critical for our experimental design, the motion contained within the targets caused the endpoints of eye movements made to those targets to be systematically biased in the direction of grating motion (motion-induced bias, MIB). We first show that this motion-induced bias can be used as a measure of sensitivity to visual motion on a trial-by-trial basis. Next, we use an exploratory approach to study whether and how the effects of reward on choice are different or similar to the effects of reward on sensitivity to visual motion measured by the MIB. In this approach, we rely on known effects of objective and subjective reward values on choice and then test those effects for the MIB.

### MIB measures sensitivity to visual motion

The motion-induced bias of a saccadic eye movement quantifies the extent to which the endpoints of saccades directed toward the drifting gratings were biased in the direction of grating motion (**Fig. 1A, Fig. 2A**). Despite the stationary position of the grating aperture, motion in the drifting sinusoid nonetheless induces a shift in the perceived position of the aperture in human subjects (De Valois & De Valois, 1991) and biases saccadic endpoints in the direction of grating drift in monkeys (Schafer & Moore, 2007). By examining the MIB in different conditions, we established that it can provide a measure of sensitivity to visual motion even when the grating motion is not behaviorally relevant.

**Figure 2.**
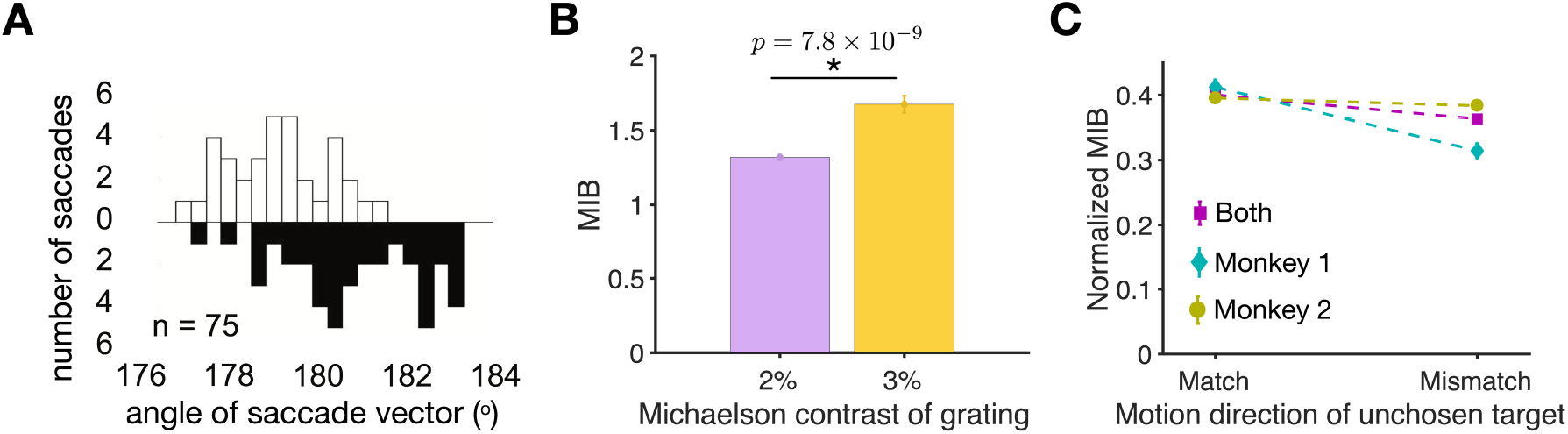
MIB measures sensitivity to visual motion. (**A**) Plotted are the example distributions of the angle of saccade vector (relative to the fixation dot) for upward (open) and downward (filled) drifting targets. (**B**) MIB significantly increased as the contrast of grating is increased from 2% (purple) to 3% (yellow). (**C**) Comparison of the z-score normalized MIB when the directions of motion in the chosen and nonchosen targets matched or did not match. The MIB is z-score normalized for each monkey separately within each session.

First, we found that the magnitude of the MIB depended on the grating contrast. More specifically, the MIB increased by 27% when the (Michaelson) contrast of grating increased from 2% to 3% (two-sided independent measures t-test, *p*=7.85 * 10^−9^; **Fig. 2B**). Second, we observed that the MIB depended almost exclusively on the motion direction of the selected target as it was only slightly affected by non-matching motion in the unchosen target (**Fig. 2C**). Specifically, the average z-score normalized MIB measured in two monkeys across all trials (mean = 0.38) was altered by only 9% when the unchosen target differed in direction of the grating motion. Together, these results demonstrate that the MIB in our task is sensitive to the properties of sensory signal (grating motion direction and contrast) and thus, can be used to measure the influence of internal factors such as subjective reward value on visual processing.

### Effects of objective reward value on choice behavior

To examine effects of global and objective reward value on monkeys’ choice behavior, we first measured how monkeys’ choice behavior tracked the target location that was globally (session-wise) more valuable, which in our task is set by reward parameter *r*. We found that target selection was sensitive to reward parameter in both monkeys and the harvested reward rate was high, averaging 0.66 and 0.65 across all sessions (including those with penalty) in monkeys 1 and 2, respectively (**Fig. 3A, B, D, E**). To better quantify monkeys’ performance, we also computed the overall harvested reward by a model that selects between the two targets with the optimal but fixed choice probability in a given session (optimal static model; see example in **Fig. 1C** in Methods) or a model in which the target with higher probability of reward was chosen on each trial (optimal dynamic model; See Methods). We found that performance of both monkeys was sub-optimal; however, the pattern of performance as a function of reward parameter for monkeys 1 and 2 resembled the behavior of the optimal static and dynamic models, respectively (**Fig. 3B, E**). Since each session of the experiment for monkey 2 was longer, we confirmed that there was no significant difference in task performance between the first and second halves of sessions for monkey 2 (difference mean±sem: 0.003 ± 0.008; two-sided paired t-test, *p* = 0.7, *d* = 0.03). Together, these results suggest that both monkeys followed the reward schedule on each session closely whereas their choice behavior was suboptimal.

**Figure 3.**
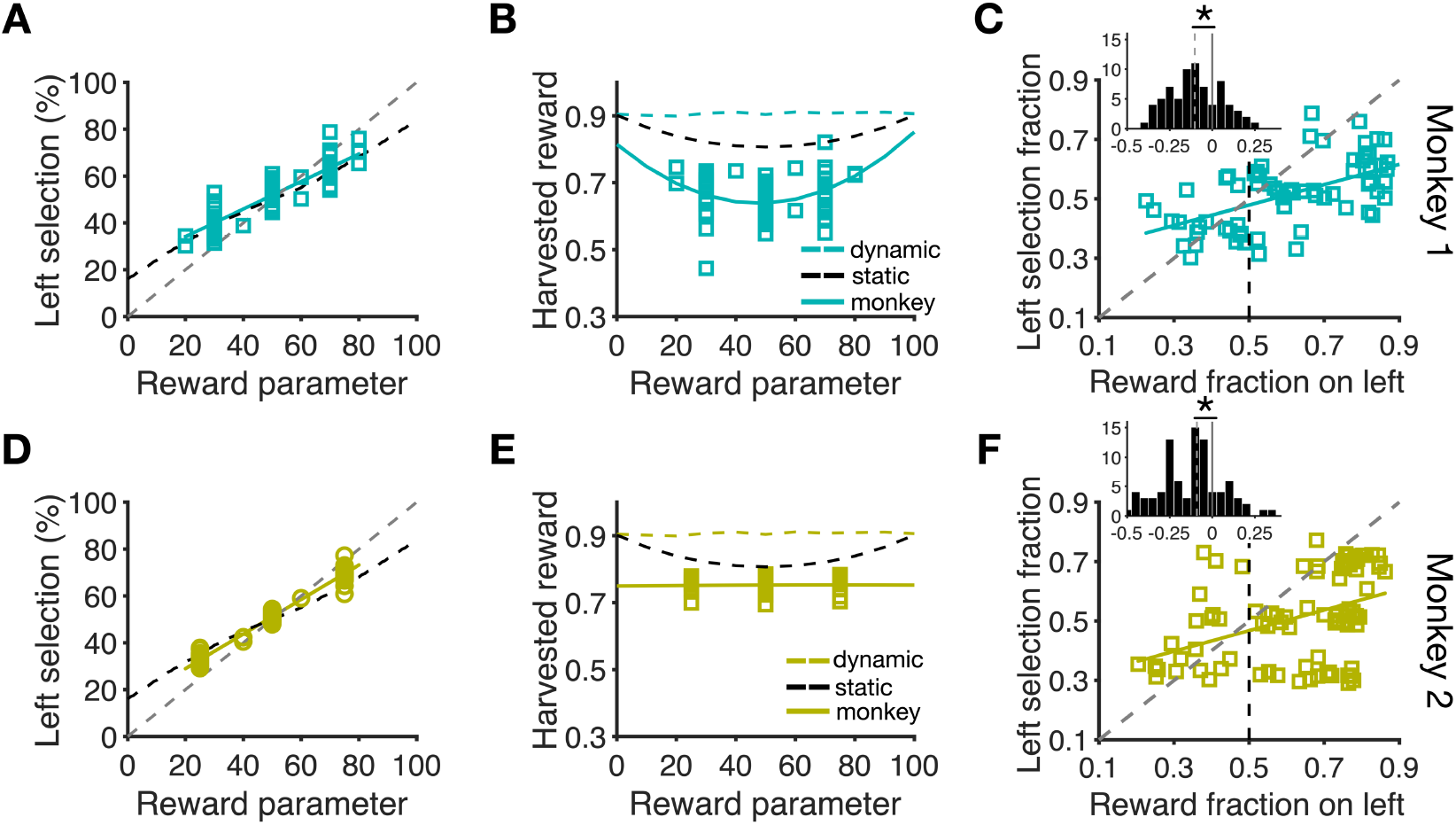
Global (session-wise) effects of reward on choice behavior. (**A**) Choice behavior was sensitive to reward parameter. Percentage of *T_L_* selections is plotted as a function of *r*, which varied across experimental sessions for monkey 1. The colored lines are linear fits, and the black dashed line shows the optimal *f*(*T_L_*) for a given value of *r* assuming selection between the two targets with a fixed probability (optimal static model). The gray dashed line shows unit slope. Each data point correspond to one session of the experiment. (**B**) The overall performance was suboptimal. Plotted is harvested rewards per trial as a function of reward parameter *r* for zero penalty sessions for monkey 1. The solid colored lines show fit using a quadratic function. The colored and black dashed lines indicate harvested reward rates of the optimal dynamic and static models, respectively. (**C**) Proportion of *T_L_* selections is plotted as a function of the fraction of harvested reward on the left target. The colored lines are linear fits and the gray dashed line shows the diagonal line corresponding to matching behavior. Monkey 1 showed significant under-matching by selecting the more rewarding target with a choice fraction smaller than reward fraction. The inset shows the difference between choice and reward fractions with negative and positive values corresponding to under- and over-matching. The gray dashed lines indicate the medians of the distributions and asterisks show the significant difference from 0 (i.e. matching) using Wilcoxon signed rank test (*p*<.05). (**D–F**) Similar to panels A-C but for monkey 2.

We also examined the global effects of reward on choice by measuring matching behavior. To that end, we compared choice and reward fractions in each session and found that both monkeys exhibited undermatching behavior (**Fig. 3C, F**). More specifically, they selected the more rewarding location with a probability that was smaller than the relative reinforcement obtained on that location (monkey 1 median(choice fraction – reward fraction) = −0.103; Wilcoxon signed rank test, *p* = 2.43 × 10^−6^, *d* = 0.65; **Fig. 3C** inset; monkey 2 median(choice fraction – reward fraction) = −0.09, *p* = 1.41 × 10 ^7^, *d* = 0.67; **Fig. 3F** inset). Furthermore, the degree of undermatching was similar for the two monkeys (diff = 0.013; Wilcoxon rank sum test, *p* = 0.58, *d* = 0.05).

### Effects of objective reward value on sensitivity to visual motion

In the previous section we observed that choice behavior is affected by objective measures of reward value in a given session. We repeated similar analyses to examine whether objective reward values have similar effects on sensitivity to visual motion measured by MIB. To that end, we first computed the correlation between the difference in the session-based average MIB for saccades to the more and less rewarding target locations and reward parameter *r* in each session. However, we did not find any evidence for such correlation for either of the two monkeys (Spearman correlation; monkey 1: *r* = 0.04, *p* = 0.8; monkey 2: *r* = 0.11, *p* = 0.39). Second, we examined whether the average MIB for all saccades in a given session was affected by the overall performance in that session. Again, we did not find any evidence for correlation between the session-based average MIB and performance for either of the two monkeys (Spearman correlation; monkey 1: *r* = −0.07, *p* = 0.54; monkey 2: *r* = 0.13, *p* = 0.21). Finally, we did a similar analysis to matching behavior to examine whether differential MIB on the two target locations is related to objective reward values of those locations. In this analysis, we computed correlation between the difference in average MIB on the better and worse target locations and the difference in total reward obtained on those locations but found no evidence for such correlation (Spearman correlation; monkey 1: *r* = 0.003, *p* = 0.99; monkey 2: *r* = 0.03, *p* = 0.83).

Together, these results indicate that unlike choice, the MIB is not affected by objective reward value of the foveated target or the overall harvested reward. Observing this dissociation, we next examined the effects of subjective reward value on choice and the MIB.

### Effects of subjective reward value on choice behavior

The analyses presented above show that the overall choice behavior was influenced by global or objective reward value of the two target locations in a given session. In contrast, sensitivity to visual motion was not affected by global or objective reward value. This difference between the influence of objective reward value on choice and MIB could simply reflect the fact that due to task design, monkeys’ choices and not MIB determine reward outcomes on current trials and influence reward probability on subsequent trials (reward probability was a function of *r* and monkeys’ choices on the preceding trials). Therefore, we next examined similarities and differences between effects of subjective reward value on choice behavior and sensitivity to visual motion.

To investigate how reward outcomes were integrated over time to estimate subjective reward values and guide monkeys’ choice behavior on each trial, we used multiple reinforcement learning (RL) models to fit the choice behavior of individual monkeys on each session of the experiment. These models assume that selection between the two targets is influenced by subjective values associated with each target, which are updated on each trial based on reward outcome (see Methods). Although reward was assigned based on the location of the two targets (left vs. right) in our experiment, the monkeys could still assume that motion direction is informative about reward. Therefore, we considered RL models in which subjective values were associated with target locations as well as RL models in which subjective values were associated with the motion of the two targets, using four different learning rules. Considering the observed undermatching behavior, we grouped learning rules depending on whether they result in the estimation of subjective value in terms of local (in time) return or income.

In RL_ret_ models, only the subjective value of the chosen target (in terms of location or motion) was updated, making them return-based models. In RL_Inc_(1) models, in addition to updating the subjective value of the chosen target, the subjective value of the unchosen target decayed at a rate similar to the subjective value of the chosen target, making these models income-based. In RL_Inc_(2) models, the subjective value of chosen and unchosen targets were allowed to decay at different rates. Finally, in RL_Inc_(3) models, we also assumed a change in the subjective value of the unchosen target or motion direction in addition to the decay. Because the subjective value of both chosen and unchosen target locations were updated on each trial in RL_Inc_(2) and RL_Inc_(3) models, we refer to these models as income-based similarly to RL_Inc_(1). However, we note that only RL_Inc_(1) models are able to estimate local income accurately.

We first compared the goodness-of-fit between the location-based and motion-based RLs using negative log likelihood (-*LL*), Akaike information criterion (AIC), and Bayesian information criterion (BIC) in order to test which of the two types of models can predict choice behavior better. Such comparisons based on the three measures yield the same results because the two types of models have the same number of parameters for a given learning rule. We found that for both monkeys, all the location-based models outperformed the motion-based RLs (**Table 1**). This demonstrates that both monkeys attributed reward outcomes to target locations more strongly than to target motions, and used subjective value attributed to target locations to perform the task.

**Table 1.** Comparison of goodness-of-fit between location-based and motion-based RL models using -LL, AIC or BIC. Δ(-LL, AIC, or BIC) shows the median of the difference between location-based and motion-based RL models fitted for each session separately. Note that all differences in goodness-of-fit measures (based on -LL, AIC, and BIC) are similar because the number of parameters is the same across location-based and motion-based models. *P*-values indicate the significance of the statistical test (two-sided sign-test) for comparing the goodness-of-fit between the location-based and motion-based RLs.

| | $RL_{ret}$ | $RL_{Inc}(1)$ | $RL_{Inc}(2)$ | $RL_{Inc}(3)$ |
| --- | --- | --- | --- | --- |
| **Monkey 1** | $\Delta(-LL, AIC, \text{ or } BIC) = -5.48$<br>$p = 2.55 * 10^{-7}$ | $\Delta(-LL, AIC, \text{ or } BIC) = -7.19$<br>$p = 2.58 * 10^{-9}$ | $\Delta(-LL, AIC, \text{ or } BIC) = -6.53$<br>$p = 1.39 * 10^{-8}$ | $\Delta(-LL, AIC, \text{ or } BIC) = -8.06$<br>$p = 5.32 * 10^{-10}$ |
| **Monkey 2** | $\Delta(-LL, AIC \text{ or } BIC) = -60.96$<br>$p = 2.74 * 10^{-21}$ | $\Delta(-LL, AIC \text{ or } BIC) = -105.71$<br>$p = 2.58 * 10^{-26}$ | $\Delta(-LL, AIC, \text{ or } BIC) = -103.46$<br>$p = 2.58 * 10^{-26}$ | $\Delta(-LL, AIC, \text{ or } BIC) = -107.25$<br>$p = 2.58 * 10^{-26}$ |

After establishing that monkeys used target location to integrate reward outcomes, we next examined how this integration was performed by comparing the quality of fit in location-based models with different learning rules. We found that for monkey 1, RL_ret_ and RL_Inc_ (1) models provided the best fit of choice data; although goodness-of-fit measures were not significantly different between these models, these models provided better fits than the RL_Inc_(2) and RL_Inc_(3) models (**Fig. 4**). Interestingly, fitting choice behavior with the RL_Inc_(1) model resulted in decay rate (α) that were close to 1 for many sessions (mean and median of α were equal to 0.77 and 1.0, respectively). This result indicates that monkey 1 integrated reward over many trials to guide its choice behavior. This is compatible with the pattern of performance as a function of reward parameter for this monkey (**Fig. 3B**), which resembles the pattern of the optimal static model.

**Figure 4.**
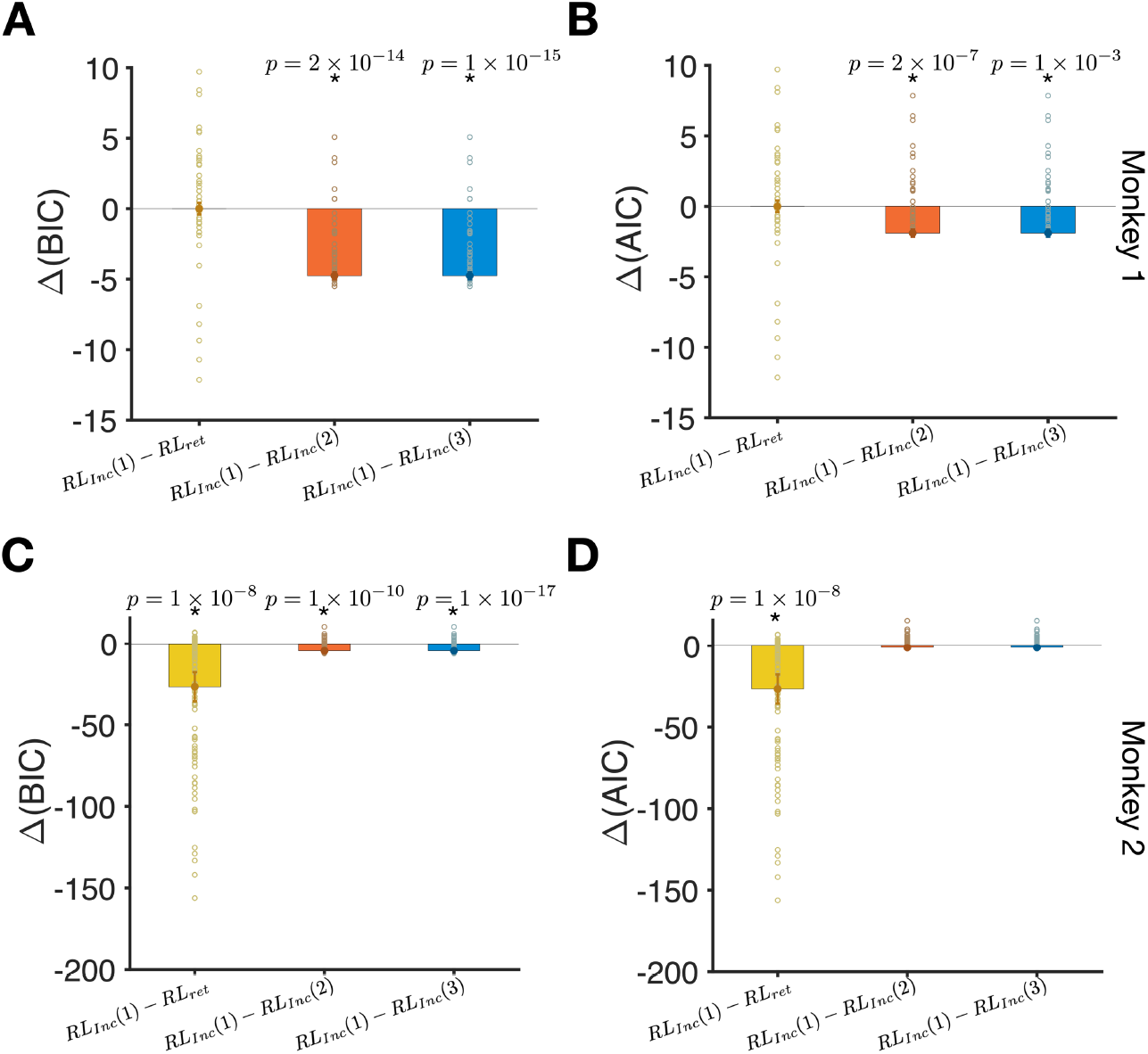
Comparison of goodness-of-fit between different location-based RL models reveals that RL_Inc_(1) model provided the best overall fit. (**A**) The difference between BIC for fits based on the RL_Inc_(1) model and the three competing models (indicated on the x-axis). Bars show the median of the difference in BIC and errors are s.e.m. Reported *p*-values are based on a twosided sign test. Each data point shows the goodness-of-fit for one session of the experiment. For monkey 1, fits based on the RL_Inc_(1) and RL_ret_ models were not significantly different. (**B**) The same as in A but based on the difference in AIC. (**C–D**) Similar to panels A and B but for monkey 2.

The same analysis for monkey 2 revealed a similar integration of reward outcomes but on a different timescale. More specifically, we found that the RL_Inc_(1) model provided the best fit for choice behavior as the goodness-of-fit in this model was better than the return-based model (RL_ret_) and more detailed income-based (RL_Inc_(2), and the RL_Inc_(3)) models (**Fig. 4**). In contrast to monkey 1, the estimated decay rate based on the RL_Inc_(1) model were much smaller than 1 for many sessions for monkey 2 (mean and median α were equal to 0.32 and 0.33, respectively). These results indicate that monkey 2 integrated reward over a shorter timescale (a few trials) than monkey 1 to guide its choice behavior. This is compatible with the pattern of performance as a function of reward parameter for this monkey (**Fig. 3E**), which resembles the pattern of the optimal dynamic model.

Together, fitting of choice behavior shows that both monkeys associated reward outcomes with the location of the chosen target. Moreover, both monkeys estimated subjective reward values in terms of income by integrating reward outcomes over multiple trials and used these values to make decisions.

### Effects of subjective reward value on sensitivity to visual motion

Our experimental design allowed us to simultaneously measure choice and the MIB, as an implicit measure of sensitivity to visual motion, on each trial. We next examined whether subjective reward value based on integration of reward outcomes over time influenced sensitivity to visual motion.

To that end, we first examined whether reward feedback had an immediate effect on the MIB in the following trial. Combining the data of the both monkeys, we found that the MIB was larger in the trials that were preceded by a rewarded rather than unrewarded trials (mean±s.e.m.: 0.03±0.009; two-sided t-test, *p* = 6.95 × 10^−4^, *d* = 0.18). When considering data from each monkey individually, however, this effect only retained significance for monkey 1 (monkey 1: mean±s.e.m.: 0.05±0.01; two-sided t-test, *p* = 6.5 × 10^−4^, *d* = 0.09; monkey 2: mean±s.e.m.: 0.01±0.01; two-sided t-test, *p* = 0.21, *d* = 0.09). These results suggest that the MIB is weakly affected by the immediate reward outcome in the preceding trial.

In the previous section, we showed that the best model for fitting choice behavior was one that estimates subjective reward value based on the income on each target location and uses the difference in incomes to drive choice behavior (RL_Inc_(1) model) (**Fig. 4**). However, it is not clear if the MIB is influenced by subjective reward values of the two targets in a similar fashion. To test this relationship, we computed correlations between the trial-by-trial MIB and estimated subjective reward values of the chosen target location, the unchosen target location, and their sum and difference. We considered subjective reward values based on both income and return (see Effects of subjective reward value on MIB in Methods).

We made several key observations. First, we found that the MIB was positively correlated with subjective reward values of both the chosen and the unchosen target (**Fig. 5A-B, Fig. 5E-F**) and as a result, was most strongly correlated with the sum of subjective reward values of the two targets (**Fig. 5C, Fig. 5G**). In contrast to choice, the MIB was poorly correlated with the difference in subjective reward values of the chosen and unchosen target (**Fig. 5D, Fig.5H, Supplementary Fig. 1**). Therefore, choice was most strongly correlated with the difference in subjective reward values, whereas the MIB was most strongly correlated with the sum of subjective reward values from the two targets. Second, although the aforementioned relationships were true for subjective reward value based on return and income, we found that correlations between the MIB and subjective return values were stronger than correlations between the MIB and subjective income values (compare **Fig. 5** and **Supplementary Fig. 2**). Third, the maximum correlation occurred for the values of τ at around 15–20 trials and for negative values of Δ_*n*_, similarly for both monkeys. This indicates that for both monkeys, the MIB was influenced by reward integrated over many trials, and the absence of reward on a given trial had a negative influence on the MIB on the following trials (Δ_*n*_ < 0).

**Figure 5.**
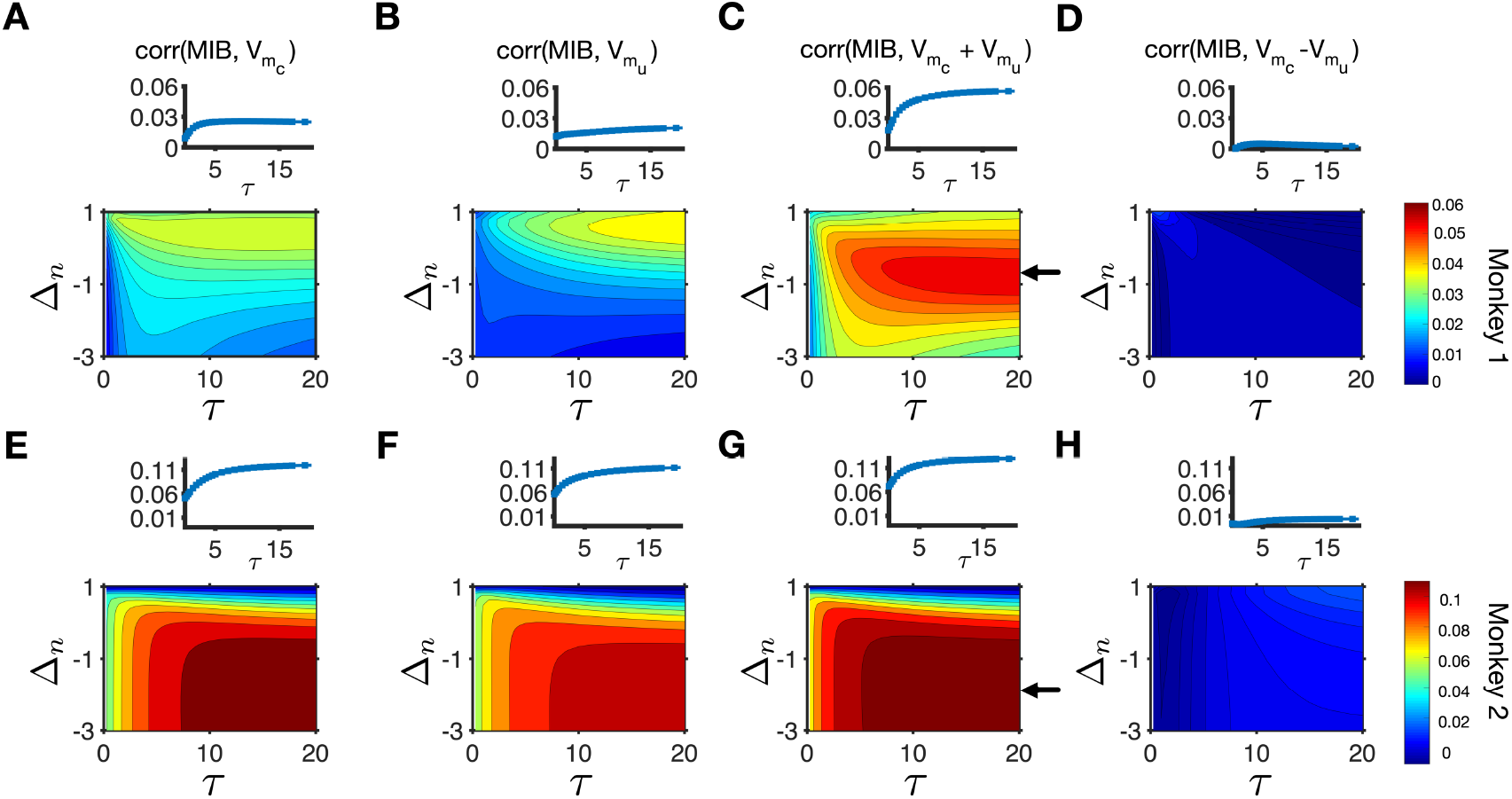
MIB was most strongly correlated with the sum of subjective reward values of the two targets based on return. (**A–D**) Plotted are the correlations between the MIB and subjective reward values of the chosen (A) and unchosen (B) targets based on return, and their sum (C) and their difference (D) for different values of τ and Δ_*n*_. The inset in each panel shows the correlation between the MIB and the corresponding subjective return values for different values of τ and a specific value of Δ_*n*_ (indicated with an arrow in the main panel C) for monkey 1. The arrow in panel C points to the value of Δ_*i*_ that results in the maximum correlation between the MIB and sum of subjective return values of the two targets for monkey 1. (**E–H**) The same as in A-D but for monkey 2. The arrow in panel G points to the value of Δ_*n*_ that results in the maximum correlation between the MIB and sum of subjective return values of the two targets for monkey 2.

Considering that local choice fraction (*f_L_* in Eq. 1) has opposite effects on *p_R_* (*T_L_*) and *p_R_* (*T_R_*) due to task design, we tested the relationship between estimated subjective values (based on return) for the two target locations. We found that the correlation between estimated reward values depends on the values of τ and Δ_*n*_ and is not always negative (data not shown). Nevertheless, for all values of τ and Δ_*n*_, the MIB was most strongly correlated with the sum of subjective reward values while being positively correlated with the value of both chosen and unchosen targets. These results indicate that the dependence of the MIB on the sum of subjective value is not driven by our specific task design.

Finally, to better illustrate distinct effects of reward on decision making and visual processing, we used two sets of parameters (τ = 15 and Δ_*n*_ = 0, τ = 15 and Δ_*n*_ = −0.5) that resulted in significant correlations between choice and targets’ subjective income values (**Supplementary Fig. 1**) and between the MIB and targets’ subjective return values in all cases (**Fig. 5**). We then used these two sets of parameters and choice history of the monkeys on the preceding trials to estimate subjective income values and return values in each trial (see Effects of subjective reward value on MIB in Methods). We then grouped trials into bins according to estimated subjective reward values of *T_L_* (left target) and *T_R_* (right target) for choice, or of the chosen and unchosen targets for the MIB, and computed the average probability of choosing the left target and the average MIB for each bin. We found that the probability of choosing the left target for both monkeys was largely determined by the difference in subjective values of the left and right targets, as can be seen from contours being parallel to the diagonals (**Fig. 6A, B, E, F**). In contrast, the MIB was largely determined by the sum of subjective values, as can be seen from contours being parallel to the second diagonals (**Fig. 6C, D, G, H)**. These results clearly demonstrate that reward has distinct effects on choice behavior and sensitivity to visual motion.

**Figure 6.**
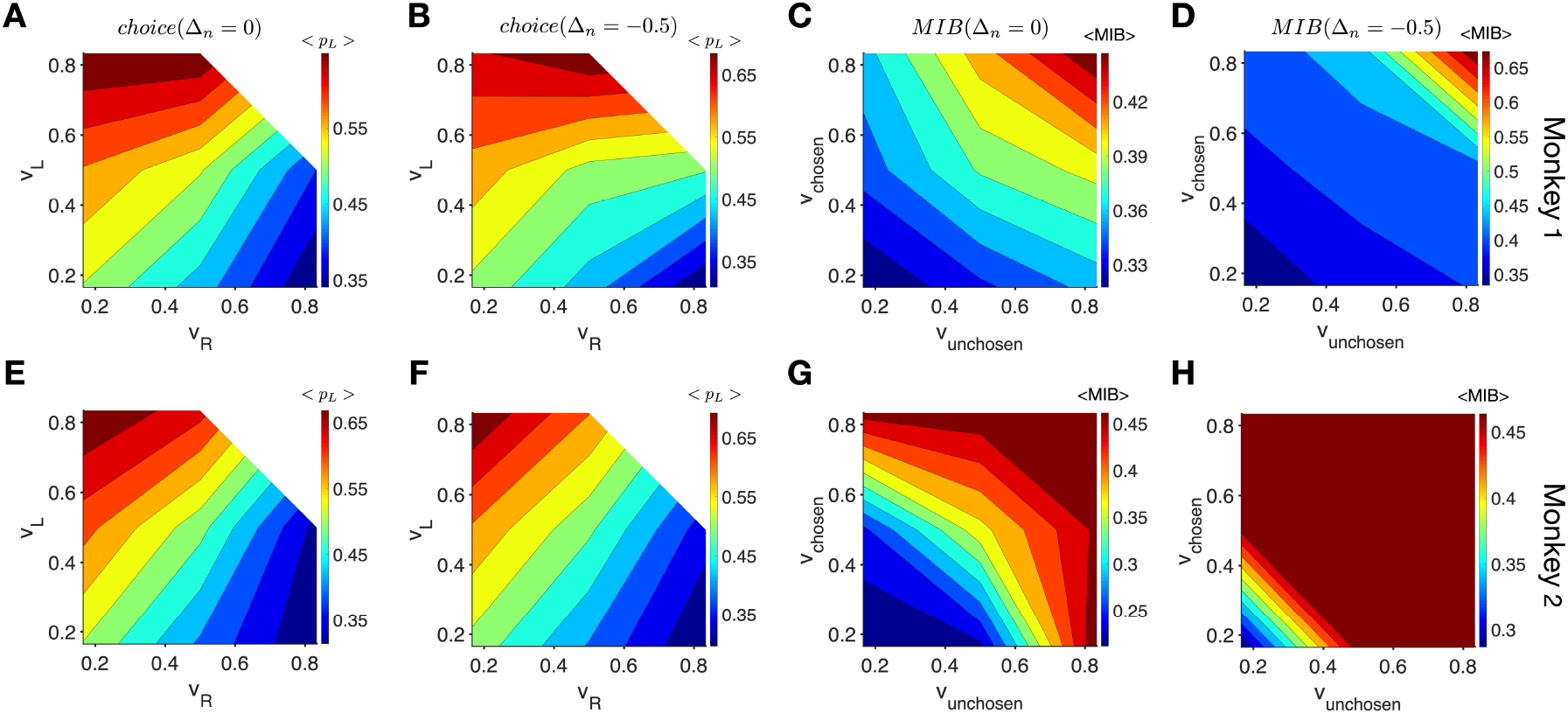
The choice probability for both monkeys was largely determined by the difference in estimated subjective values whereas the MIB was largely determined by the sum of subjective values of targets. (**A–B**) Plots show the probability of choosing the left target as a function of subjective values of the left and right targets for monkey 1, using τ = 15 and two values of Δ_*n*_ as indicated on the top. (**C–D**) Plots show the MIB as a function of subjective values of the chosen and unchosen targets for monkey 1, using τ = 15 and two values of Δ_*n*_ as indicated on the top. (**E–H**) The same as in A-D but for monkey 2.

## Discussion

Experimental paradigms with dynamic reward schedules have been extensively used in different animal models to study how reward shapes choice behavior on a trial-by-trial basis (Barraclough et al., 2004; Donahue & Lee, 2015; Herrnstein, Loewenstein, Prelec, & Vaughan, 1993; Lau & Glimcher, 2005; Li, McClure, King-Casas, & Montague, 2006; Sugrue et al., 2004). A general finding is that animals integrate reward outcomes on one or more timescales in order to estimate subjective reward value and determine choice. In contrast, the influence of reward on selective processing of visual information, which is often described as attentional deployment, has been mainly studied using fixed reward schedules with unequal reward outcomes (B. A. Anderson et al., 2011a, 2011b; Barbaro et al., 2017; Della Libera & Chelazzi, 2006, 2009; Hickey et al., 2010, 2014; Hickey & Peelen, 2017; Peck et al., 2009). The main findings from these studies are that targets or features associated with larger reward can more strongly capture attention and alter visual processing immediately or even after extended periods of time (reviwed in B. A. Anderson, 2013, 2016).

However, it has proven difficult to link the effects of reward on saccadic choice and selective processing of visual information mainly because of separate measurements of these effects in different tasks. Indeed, the poorly described relationship between reward expectation and the processing of visual information has been implicated as a confounding factor in the interpretation of many past behavioral and neurophysiological results (Maunsell, 2004, 2015). An exception to this is a study by Serences (2008) in which the author utilized a task with dynamic reward schedule to demonstrate that the activity in visual cortex is modulated by reward history (i.e., integrated reward outcomes over many trials). Compatible with these results, we find that processing of visual information is affected by subjective reward value estimated by integration of reward outcomes over many trials.

Using tasks designed specifically to dissociate subjective reward value from a target’s behavioral significance, or salience, a few studies have identified brain areas that respond primarily to the expected reward or the salience of a target (or both) in various species including rats (Lin & Nicolelis, 2008), monkeys (Roesch & Olson, 2004), and humans (Anderson et al., 2003; Cooper & Knutson, 2008; Jensen et al., 2007; Litt, Plassmann, Shiv, & Rangel, 2011). However, in these studies, the saliency signal observed in neural responses might reflect a number of different processes, such as motivation, attention, motor preparation, or some combination of these. In the present work, we exploited the influence of visual motion on saccades as an independent and implicit measure of visual processing during value-based decision making. This enabled us for the first time to measure choice and visual processing simultaneously and to test whether reward has differential effects on these two processes.

Although motion was not predictive of reward and thus processing of motion direction was not required to obtain a reward, we found that similar to decision making, visual processing was influenced by subjective reward values of the two targets. However, subjective reward values of the two targets affected visual processing differently than how they affected choice in three ways. First, although choice was correlated most strongly with the difference between subjective values of chosen and unchosen targets, visual processing was most strongly correlated with the sum of subjective values of the two targets. The latter indicates that the overall subjective value of targets in a given environment could influence the quality of sensory processing in that environment. Second, choice was more strongly affected by the subjective income value of the target whereas sensitivity to visual motion was more strongly affected by subjective return values of the targets. Third, the time constant of reward integration, and the impact of no-reward were different between decision making and visual information processing. In contrast to subjective reward value, we found that objective reward value only affected choice and not sensitivity to visual motion. Together, these results point to multiple systems for reward integration in the brain.

We found certain differences between the results for the two monkeys that could indicate that they used different, idiosyncratic strategies for performing the task. For example, fitting results of reinforcement learning models indicated that monkey 1 used the reward history over many trials to direct its choice behavior. In contrast, monkey 2 used the reward history over few trials to direct its choice behavior. This difference was also apparent in the correlation between choice and the difference in subjective values of the two target locations. Despite this difference in integration time constant, choice in both monkeys was most strongly correlated with the difference between estimated subjective values of the two targets. Furthermore, the MIB for both monkeys was most strongly correlated with the sum of estimated subjective values of the two targets, even though they integrated reward outcomes on different timescales.

The observed differences in reward effects on visual processing and decision making have important implications for the involved brain structures and underlying neural mechanisms. First, they suggest that brain structures involved in decision making and processing of visual information receive distinct sets of value-based input; e.g., ones that integrate reward over a different number of trials. The set of input affecting decision making carries information about subjective reward value of individual targets whereas the set that affects visual processing carries information about the sum of subjective values of targets. Indeed, there are more neurons in the anterior cingulate cortex and other prefrontal areas that encode the sum of subjective value of available options than subjective value of a given option (Kim, Hwang, Seo, & Lee, 2009), and these neurons might contribute to enhanced sensory processing. In addition, it has been shown that the activity of basal forebrain neurons increases with the sum of subjective values of choice array options (Ledbetter, Chen, & Monosov, 2016) and this could enable basal forebrain to guide visual processing and attention based on reward feedback independently of how reward controls choice behavior (Monosov, 2020). Finally, the frontal eye field (FEF) also receives inputs from the supplementary eye field (SEF), which contains neurons whose activity reflects subjective reward value of the upcoming saccade (Chen & Stuphorn, 2015). Such input from the SEF could drive target selection in the FEF. Importantly, our findings can be used in future experiments to tease apart neural substrates by which reward influences visual processing and decision making.

Second, a plausible mechanism that could contribute to the observed differences in the effects of reward is the differential influence of dopaminergic signaling on the functions of FEF neurons. Recent work demonstrates that the modulatory influence of the FEF on sensory activity within visual cortex is mediated principally by D1 receptors, and that D2-mediated activity is not involved (Noudoost & Moore, 2011). However, activity mediated through both receptor subtypes contributes to target selection, albeit in different ways (Noudoost & Moore, 2011; Soltani, Noudoost, & Moore, 2013). This evidence indicates that the neural mechanisms underlying target selection and visual processing are separable if only in terms of the involvement of different dopaminergic signals. Considering the known role of dopamine in reward processing (Schultz, 2007) and synaptic plasticity (Calabresi, Picconi, Tozzi, & Di Filippo, 2007), these two dopaminergic signaling pathways may provide a mechanism for the separate effects of reward on sensory processing and selection.

Third, in most choice tasks with dynamic reward schedules, local subjective return and income values are typically correlated, and the question of which quantity is the critical determinant of behavior has been debated for many years (Corrado et al., 2005; Gallistel & Gibbon, 2000; Gallistel, Mark, King, & Latham, 2001; Herrnstein & Prelec, 1991; Mark & Gallistel, 1994; Soltani & Wang, 2006). The observation that differences in subjective income values are a better predictor of choice behavior may reflect the fact that income values provide information about which target is globally more valuable in each session of the task. In contrast, the dependence of visual processing on the sum of subjective return values is more unexpected. This indicates that visual processing may more strongly depend on target-specific reward integration because the return value of a given target is updated only after selection of that target.

Finally, the separable influences of reward could be crucial for flexible behavior required in dynamic and high-dimensional reward environments (Farashahi, Rowe, et al., 2017). For example, processing of visual information of the saccade target that has multiple visual features based on the sum of subjective reward values of available targets could allow processing of previously neglected information from the less rewarding targets and thus, improve exploration. Future studies are needed to test whether disruption of this processing can reduce flexibility in target selection and choice behavior.

## Acknowledgments

We thank Vince McGinty for helpful comments on an earlier version of this manuscript. We also thank D.S. Aldrich for technical assistance. This work was supported by NIH Grant EY014924 (TM), NIH Grant DA047870 (AS), NSF EPSCoR Award #1632738 (AS), an NDSEG fellowship (RJS), and predoctoral NRSA fellowship F31MH078490 (RJS).

## Supplementary Figures

**Supplementary Figure 1.**
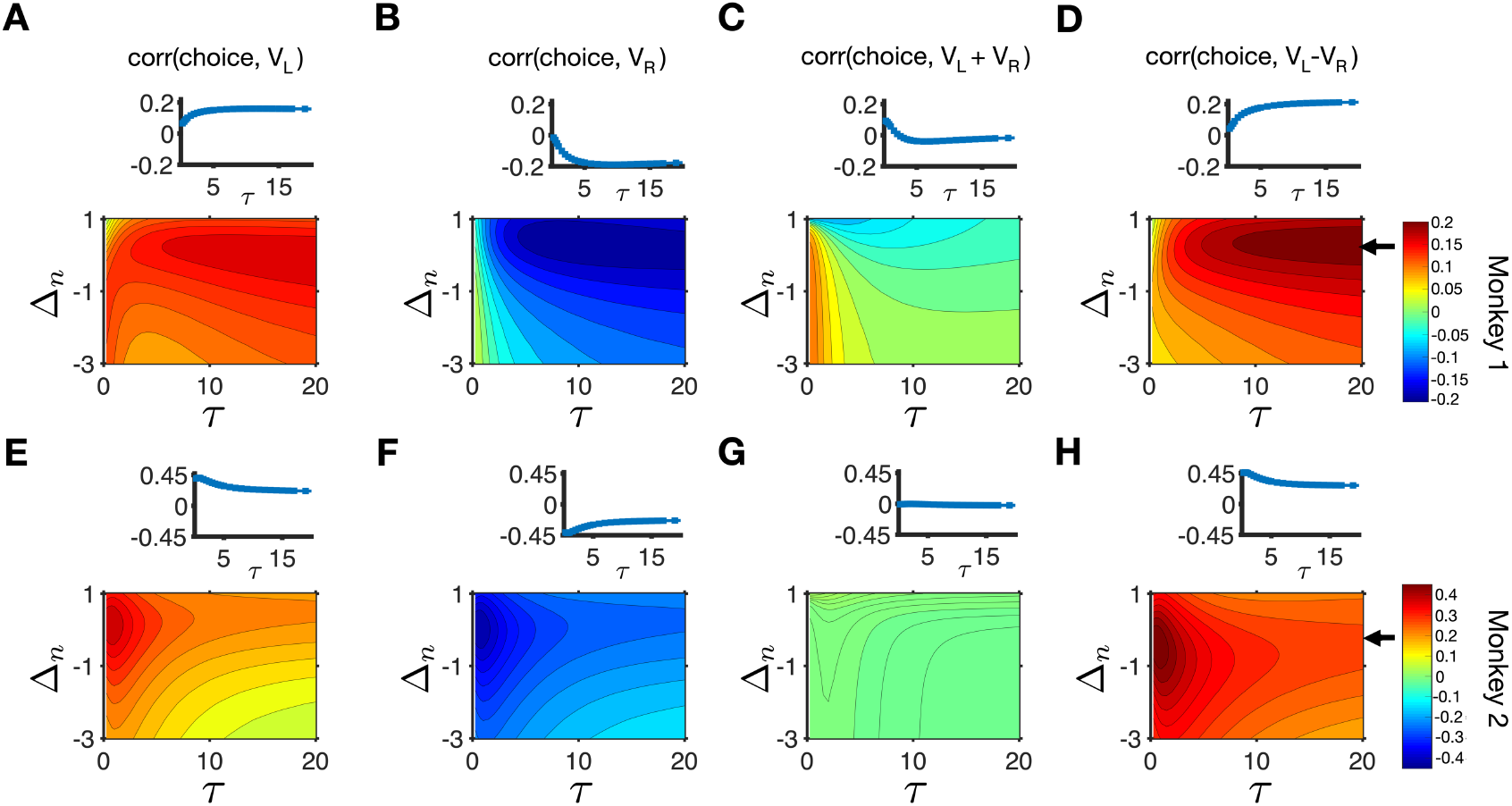
Choice was mainly correlated with the difference in subjective values of the two targets in terms of income. (**A–D**) Plotted are the correlations between selection of the left target and subjective income values of the left (A) and right (B) targets, and their sum (C) and their difference (D) for different values of τ and a specific value of Δ_*n*_ (indicated with an arrow in the inset) for monkey 1. The inset in each panel shows the correlation between choice and the corresponding subjective income values of targets for different values of τ and Δ_*n*_. The arrow in panel D points to the value of Δ_*n*_ that results in the maximum correlation between selection of the left target and the difference in subjective income values of the two targets for monkey 1. (**E–H**) The same as in A-D but for monkey 2. The arrow in panel H points to the value of Δ_*n*_ that results in the maximum correlation between selection of the left target and the difference in subjective income values of the two targets for monkey 2.

**Supplementary Figure 2.**
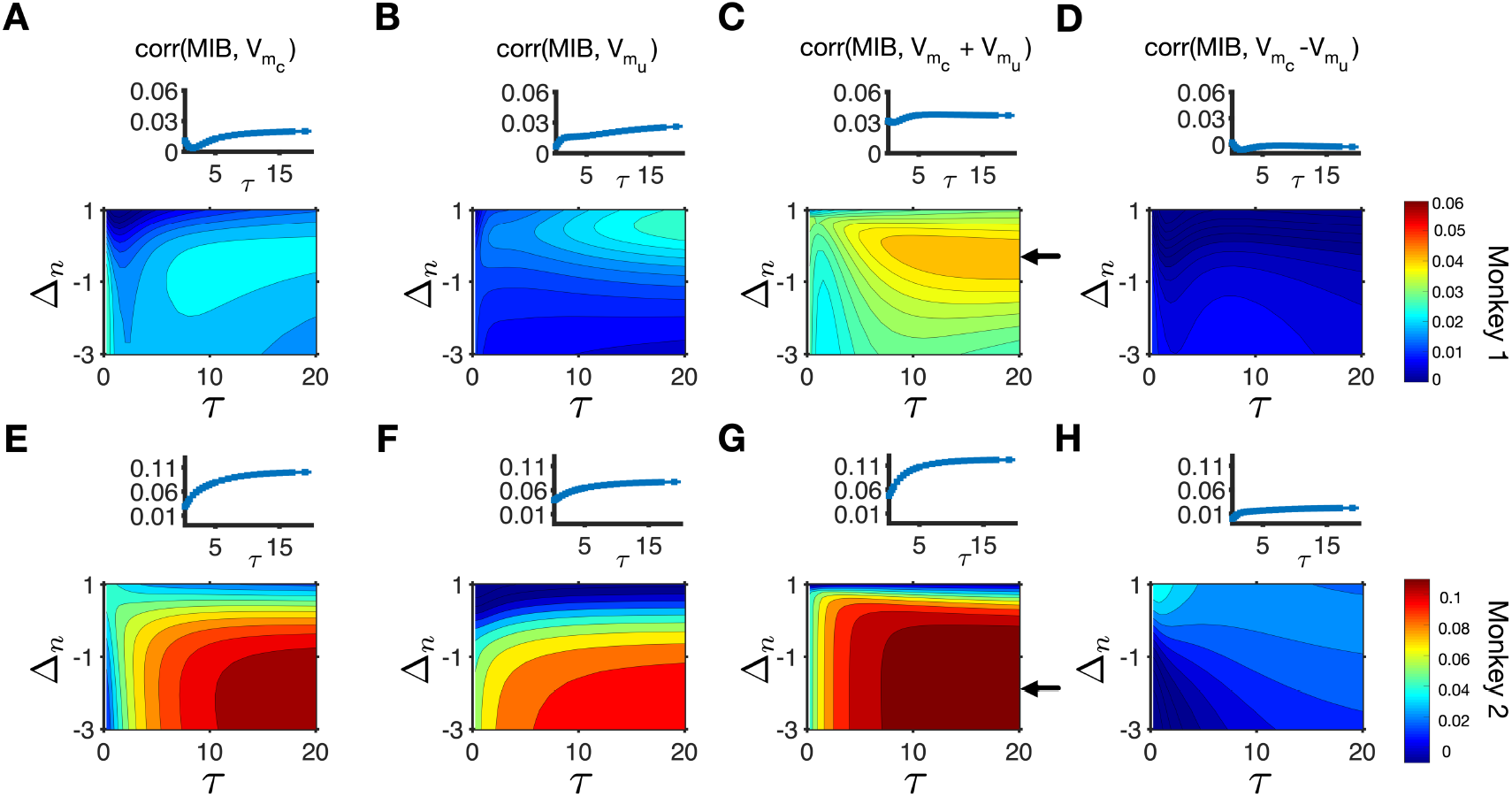
Correlation between the MIB and subjective income of the chosen and unchosen targets. (**A–D**) Plotted are the correlations between the MIB and subjective income values of the chosen (A) and unchosen (B) targets, and their sum (C) and their difference (D) for different values of τ and a specific value of Δ_*n*_ (indicated with an arrow in the inset) for monkey 1. The inset in each panel shows the correlation between the MIB and the corresponding subjective values (in terms of income) for different values of τ and Δ_*n*_. The arrow in panel C points to the value of Δ_*n*_ that results in the maximum correlation between the MIB and the sum of subjective income values of the two targets for monkey 1. (**E**–**H**) The same as in A-D but for monkey 2. The arrow in panel G points to the value of Δ_*n*_ that results in the maximum correlation between the MIB and the sum of subjective income values of the two targets for monkey 2.

## Notes

**Conflict of interest:** The authors declare no competing interests.

### Competing Interest Statement

The authors have declared no competing interest.

### Summary of Updates

This revision provides additional information about Methods and motivation behind the study.

## References

Abe, N., & Takeuchi, J. (1993). The “Lob-Pass” problem and an on-line learning model of rational choice.” In Proc. 6th Ann. Conf. on Comp. Learning Theory (pp. 422–428). Retrieved from https://doi.org/10.1145/168304.168389

Anderson, A. K., Christoff, K., Stappen, I., Panitz, D., Ghahremani, D. G., Glover, G., … Sobel, N. (2003). Dissociated neural representations of intensity and valence in human olfaction. Nature Neuroscience, 6(2), 196–202. https://doi.org/10.1038/nn1001

Anderson, B. A. (2013). A value-driven mechanism of attentional selection. Journal of Vision, 13(3), 7–7. https://doi.org/10.1167/13.3.7

Anderson, B. A. (2016). The attention habit: How reward learning shapes attentional selection. Annals of the New York Academy of Sciences, 1369(1). https://doi.org/10.1111/nyas.12957

Anderson, B. A., Laurent, P. A. P. A., & Yantis, S. (2011a). Value-driven attentional capture. Proceedings of the National Academy of Sciences, 108(25), 10367–10371. https://doi.org/10.1073/pnas.1104047108

Anderson, B. A., Laurent, P. A., & Yantis, S. (2011b). Learned value magnifies salience-based attentional capture. PLoS ONE, 6(11), e27926. https://doi.org/10.1371/journal.pone.0027926

Barbaro, L., Peelen, M. V, & Hickey, C. (2017). Valence, not utility, underlies reward-driven prioritization in human vision. The Journal of Neuroscience, 37(43), 1128–17. https://doi.org/10.1523/JNEUROSCI.1128-17.2017

Bari, B. A., Grossman, C. D., Lubin, E. E., Rajagopalan, A. E., Cressy, J. I., & Cohen, J. Y. (2019). Stable Representations of Decision Variables for Flexible Behavior. Neuron, 103(5), 922–933.e7. https://doi.org/10.1016/j.neuron.2019.06.001

Barraclough, D. J., Conroy, M. L., & Lee, D. (2004). Prefrontal cortex and decision making in a mixed-strategy game. Nature Neuroscience, 7(4), 404–410. https://doi.org/10.1038/nn1209

Calabresi, P., Picconi, B., Tozzi, A., & Di Filippo, M. (2007). Dopamine-mediated regulation of corticostriatal synaptic plasticity. Trends Neurosci, 30(5), 211–219. https://doi.org/10.1016/j.tins.2007.03.001

Chen, X., & Stuphorn, V. (2015). Sequential selection of economic good and action in medial frontal cortex of macaques during value-based decisions. ELife, 4, 1–24. https://doi.org/10.7554/eLife.09418

Cooper, J. C., & Knutson, B. (2008). Valence and salience contribute to nucleus accumbens activation. NeuroImage, 39(1), 538–547. Retrieved from 10.1016/j.neuroimage.2007.08.009

Corrado, G. S., Sugrue, L. P., Seung, H. S., & Newsome, W. T. (2005). Linear-Nonlinear-Poisson Models of Primate Choice Dynamics. Journal of the Experimental Analysis of Behavior, 84(3), 581–617. https://doi.org/10.1901/jeab.2005.23-05

Costa, V. D., Dal Monte, O., Lucas, D. R., Murray, E. A., & Averbeck, B. B. (2016). Amygdala and Ventral Striatum Make Distinct Contributions to Reinforcement Learning. Neuron, 92(2), 505–517. https://doi.org/10.1016/j.neuron.2016.09.025

De Valois, R. L., & De Valois, K. K. (1991). Vernier acuity with stationary moving Gabors. Vision Research, 31(9), 1619–1626. https://doi.org/10.1016/0042-6989(91)90138-U

Della Libera, C., & Chelazzi, L. (2006). Visual selective attention and the effects of monetary rewards. Psychological Science, 17(3), 222–227. Retrieved from 10.1111/j.1467-9280.2006.01689.x

Della Libera, C., & Chelazzi, L. (2009). Learning to attend and to ignore is a matter of gains and losses. Psychological Science, 20(6), 778–784. https://doi.org/10.1111/j.1467-9280.2009.02360.x

Donahue, C. H., & Lee, D. (2015). Dynamic routing of task-relevant signals for decision making in dorsolateral prefrontal cortex. Nature Neuroscience, 18(2), 295–301. https://doi.org/10.1038/nn.3918

Farashahi, S., Azab, H., Hayden, B., & Soltani, A. (2018). On the flexibility of basic risk attitudes in monkeys. The Journal of Neuroscience, 38(18), 4383–4398. https://doi.org/10.1523/JNEUROSCI.2260-17.2018

Farashahi, S., Donahue, C. H., Khorsand, P., Seo, H., Lee, D., & Soltani, A. (2017). Metaplasticity as a Neural Substrate for Adaptive Learning and Choice under Uncertainty. Neuron, 94(2), 401–414.e6. https://doi.org/10.1016/j.neuron.2017.03.044

Farashahi, S., Rowe, K., Aslami, Z., Lee, D., & Soltani, A. (2017). Feature-based learning improves adaptability without compromising precision. Nature Communications, 8(1). https://doi.org/10.1038/s41467-017-01874-w

Fuchs, A. F., & Robinson, D. A. (1966). A method for measuring horizontal and vertical eye movement chronically in the monkey. Journal of Applied Physiology, 21(3), 1068–1070. Retrieved from 10.1152/jappl.1966.21.3.1068

Gallistel, C. R., & Gibbon, J. (2000). Time, rate, and conditioning. Psychological Review, 107(2), 289–344. Retrieved from 10.1037/0033-295x.107.2.289

Gallistel, C. R., Mark, T. A., King, A. P., & Latham, P. E. (2001). The rat approximates an ideal detector of changes in rates of reward: implications for the law of effect. Journal of Experimental Psychology Animal Behavior Processes, 27(4), 354–372. Retrieved from 10.1037//0097-7403.27.4.354

Glimcher, P. W. (2003). The neurobiology of visual-saccadic decision making. Annual Review of Neuroscience, 26(1), 133–179. https://doi.org/10.1146/annurev.neuro.26.010302.081134

Herrnstein, R. J. (1961). Relative and absolute strength of response as a function of frequency of reinforcement. Journal of the Experimental Analysis of Behavior, 4(3), 267–272. https://doi.org/10.1901/jeab.1961.4-267

Herrnstein, R. J., Loewenstein, G. F., Prelec, D., & Vaughan, W. (1993). Utility Maximization and Melioration: Internalities in Individual Choice. Journal of Behavioral Decision Making, 6(3), 149–185. Retrieved from 10.1002/bdm.3960060302

Herrnstein, R. J., & Prelec, D. (1991). Melioration: a theory of distributed choice. Journal of Economic Perspectives, 5(3), 137–156. https://doi.org/10.1257/jep.5.3.137

Hickey, C., Chelazzi, L., & Theeuwes, J. (2010). Reward changes salience in human vision via the anterior cingulate. Journal of Neuroscience, 30(33), 11096–11103. https://doi.org/10.1523/JNEUROSCI.1026-10.2010

Hickey, C., Chelazzi, L., & Theeuwes, J. (2014). Reward-Priming of Location in Visual Search. PLoS ONE, 9(7), e103372. https://doi.org/10.1371/journal.pone.0103372

Hickey, C., & Peelen, M. V. (2017). Reward Selectively Modulates the Lingering Neural Representation of Recently Attended Objects in Natural Scenes. The Journal of Neuroscience, 37(31), 7297–7304. https://doi.org/10.1523/JNEUROSCI.0684-17.2017

Hikosaka, O. (2007). Basal ganglia mechanisms of reward-oriented eye movement. Annals of the New York Academy of Sciences, 1104(1), 229–249. https://doi.org/10.1196/annals.1390.012

Itti, L., & Koch, C. (2000). A saliency-based search mechanism for overt and covert shifts of visual attention. Vision Research, 40(10–12), 1489–1506. https://doi.org/10.1016/S0042-6989(99)00163-7

Jensen, J., Smith, A., Willeit, M., Crawley, A., Mikulis, D., Vitcu, I., & Kapur, S. (2007). Separate brain regions code for salience vs. valence during reward prediction in humans. Human Brain Mapping, 28(4), 294–302. https://doi.org/10.1002/hbm.20274

Judge, S. J., Richmond, B. J., & Chu, F. C. (1980). Implantation of magnetic search coils for measurement of eye position: an improved method. Vision Research, 20(6), 535–538. Retrieved from 10.1016/0042-6989(80)90128-5

Kim, S., Hwang, J., Seo, H., & Lee, D. (2009). Valuation of uncertain and delayed rewards in primate prefrontal cortex. Neural Networks, 22(3), 294–304. https://doi.org/10.1016/J.NEUNET.2009.03.010

Lau, B., & Glimcher, P. W. (2005). Dynamic Response-by-Response Models of Matching Behavior in Rhesus Monkeys. Journal of the Experimental Analysis of Behavior, 84(3), 555–579. https://doi.org/10.1901/jeab.2005.110-04

Lau, B., & Glimcher, P. W. (2007). Action and outcome encoding in the primate caudate nucleus. Journal of Neuroscience, 27(52), 14502–14514. https://doi.org/10.1523/JNEUROSCI.3060-07.2007

Ledbetter, N. M., Chen, C. D., & Monosov, I. E. (2016). Multiple mechanisms for processing reward uncertainty in the primate basal forebrain. Journal of Neuroscience, 36(30), 7852–7864. https://doi.org/10.1523/JNEUROSCI.1123-16.2016

Li, J., McClure, S. M., King-Casas, B., & Montague, P. R. (2006). Policy adjustment in a dynamic economic game. PLoS ONE, 1(1). https://doi.org/10.1371/journal.pone.0000103

Lin, S. C., & Nicolelis, M. A. (2008). Neuronal ensemble bursting in the basal forebrain encodes salience irrespective of valence. Neuron, 59(1), 138–149. https://doi.org/10.1016/j.neuron.2008.04.031

Liston, D. B., & Stone, L. S. (2008). Effects of prior information and reward on oculomotor and perceptual choices. Journal of Neuroscience, 28(51), 13866–13875. https://doi.org/10.1523/JNEUROSCI.3120-08.2008

Litt, A., Plassmann, H., Shiv, B., & Rangel, A. (2011). Dissociating valuation and saliency signals during decision-making. Cerebral Cortex, 21(1), 95–102. https://doi.org/10.1093/cercor/bhq065

Mark, T. A., & Gallistel, C. R. (1994). Kinetics of matching. Journal of Experimental Psychology Animal Behavior Processes, 20(1), 79–95. Retrieved from 10.1037/0097-7403.20.1.79

Markowitz, D. A., Shewcraft, R. A., Wong, Y. T., & Pesaran, B. (2011). Competition for Visual Selection in the Oculomotor System. Journal of Neuroscience, 31(25), 9298–9306. https://doi.org/10.1523/JNEUROSCI.0908-11.2011

Maunsell, J. H. R. (2004). Neuronal representations of cognitive state: reward or attention? Trends in Cognitive Sciences, 8(6), 261–265. https://doi.org/10.1016/j.tics.2004.04.003

Maunsell, J. H. R. (2015). Neuronal mechanisms of visual attention. Annual Review of Vision Science, 1(1), 373–391. https://doi.org/10.1146/annurev-vision-082114-035431

Monosov, I. E. (2020). How Outcome Uncertainty Mediates Attention, Learning, and Decision-Making. Trends in Neurosciences, 43(10), 795–809. https://doi.org/10.1016/j.tins.2020.06.009

Moore, T., & Fallah, M. (2001). Control of eye movements and spatial attention. Proceedings of the National Academy of Sciences, 98(3), 1273–1276. https://doi.org/10.1073/pnas.021549498

Navalpakkam, V., Koch, C., Rangel, A., & Perona, P. (2010). Optimal reward harvesting in complex perceptual environments. Proceedings of the National Academy of Sciences, 107(11), 5232–5237. https://doi.org/10.1073/pnas.0911972107

Noudoost, B., & Moore, T. (2011). Control of visual cortical signals by prefrontal dopamine. Nature, 474(7351), 372–375. https://doi.org/10.1038/nature09995

Peck, C. J., Jangraw, D. C., Suzuki, M., Efem, R., & Gottlieb, J. (2009). Reward modulates attention independently of action value in posterior parietal cortex. Journal of Neuroscience, 29(36), 11182–11191. https://doi.org/10.1523/JNEUROSCI.1929-09.2009

Platt, M. L., & Glimcher, P. W. (1999). Neural correlates of decision variables in parietal cortex. Nature, 400(6741), 233–238. https://doi.org/10.1038/22268

Rakhshan, M., Lee, V., Chu, E., Harris, L., Laiks, L., Khorsand, P., & Soltani, A. (2020). Influence of Expected Reward on Temporal Order Judgment. Journal of Cognitive Neuroscience, 32(4), 674–690. https://doi.org/10.1162/jocn_a_01516

Roesch, M. R., & Olson, C. R. (2004). Neuronal activity related to reward value and motivation in primate frontal cortex. Science, 304(5668), 307–310. https://doi.org/10.1126/science.1093223

Schafer, R. J., & Moore, T. (2007). Attention governs action in the primate frontal eye field. Neuron, 56(3), 541–551. https://doi.org/10.1016/j.neuron.2007.09.029

Schultz, W. (2007). Multiple dopamine functions at different time courses. Annual Review of Neuroscience, 30, 259–288. Retrieved from 10.1146/annurev.neuro.28.061604.135722

Schütz, A. C., Trommershäuser, J., & Gegenfurtner, K. R. (2012). Dynamic integration of information about salience and value for saccadic eye movements. Proceedings of the National Academy of Sciences of the United States of America, 109(19), 7547–7552. https://doi.org/10.1073/pnas.1115638109

Soltani, A., Noudoost, B., & Moore, T. (2013). Dissociable dopaminergic control of saccadic target selection and its implications for reward modulation. Proceedings of the National Academy of Sciences, 110(9), 3579–3584. https://doi.org/10.1073/pnas.1221236110

Soltani, A., & Wang, X.-J. X.-J. (2006). A Biophysically Based Neural Model of Matching Law Behavior: Melioration by Stochastic Synapses. Journal of Neuroscience, 26(14), 3731–44. Retrieved from http://www.jneurosci.org/content/26/14/3731.short

Soltani, A., & Wang, X. J. (2008). From biophysics to cognition: reward-dependent adaptive choice behavior. Current Opinion in Neurobiology, 18(2), 209–216. https://doi.org/10.1016/j.conb.2008.07.003

Squire, R. F., Noudoost, B., Schafer, R. J., & Moore, T. (2013). Prefrontal contributions to visual selective attention. Annual Review of Neuroscience, 36(1), 451–466. https://doi.org/10.1146/annurev-neuro-062111-150439

Strait, C. E., Blanchard, T. C., & Hayden, B. Y. (2014). Reward value comparison via mutual inhibition in ventromedial prefrontal cortex. Neuron, 82(6), 1357–1366. https://doi.org/10.1016/j.neuron.2014.04.032

Sugrue, L. P., Corrado, G. S., & Newsome, W. T. (2004). Matching behavior and the representation of value in the parietal cortex. Science, 304(5678), 1782–1787. https://doi.org/10.1126/science.1094765

Sugrue, L. P., Corrado, G. S., & Newsome, W. T. (2005). Choosing the greater of two goods: Neural currencies for valuation and decision making. Nature Reviews Neuroscience, 6(5), 363–375. https://doi.org/10.1038/nrn1666

Volkow, N. D., Wang, G. J., Kollins, S. H., Wigal, T. L., Newcorn, J. H., Telang, F., … Swanson, J. M. (2009). Evaluating dopamine reward pathway in ADHD: clinical implications. JAMA -Journal of the American Medical Association, 302(10), 1084–1091. https://doi.org/10.1001/jama.2009.1308

